# Evaluation of Data-based Motion Correction Techniques for High Temporal Resolution Functional PET

**DOI:** 10.1101/2025.08.27.672614

**Authors:** Pia Falb, Murray B. Reed, Sebastian Klug, Matej Murgaš, Godber M. Godbersen, Clemens Schmidt, Lukas Nics, Marcus Hacker, Rupert Lanzenberger, Andreas Hahn

## Abstract

Functional Positron Emission Tomography (fPET) data offers novel insights into brain energy demands and molecular connectivity. Recent advances in improving temporal resolutions for this imaging technique have opened up new research possibilities. However, lower signal-to-noise ratios (SNR) inherent to short PET frames bring into question whether current realignment approaches still provide appropriate motion correction. Thus, we aimed to evaluate the effectiveness of standard motion correction methods and explore potential improvements for high temporal resolution fPET with 3s frames. We investigated two techniques aimed at improving the SNR to facilitate more accurate realignment of fPET images, in comparison to conventional motion correction: a deep-learning technique based on the application of a conditional generative adversarial network and an exponentially weighted sliding window average. Performance was evaluated by correlating rigid motion parameters between approaches and with simultaneously acquired fMRI data, and by assessing magnitudes of task-induced activation. Our results indicate that neither of the two methods substantially improve mitigation of motion artefacts. Given the increased computational effort of both techniques, we propose that the standard motion correction procedure is adequate for processing high temporal resolution fPET data. Nevertheless, future development of targeted strategies to enhance motion correction may further advance this imaging technique.

## 1. Introduction

Correction of head motion in neuroimaging data, acquired with positron emission tomography (PET) and functional magnetic resonance imaging (MRI) has been a common problem for researchers. Movement can substantially deteriorate the image quality and introduce artefacts, which might distort the information that is obtained from the scans. This is particularly relevant for clinical research(1,2), as a considerable portion of clinical populations is especially prone to movement. In particular, patients suffering from certain neurological disorders, such as Alzheimer’s (3) or Parkinson’s disease (4), tend to move more often. Thus, techniques for appropriate motion correction are essential to increase the reliability of research findings and the respective conclusions.

Several algorithms are available to correct for motion-induced artefacts in dynamic neuroimaging data, which are largely the result of rigid head movement. These corrective procedures include MRI-based motion correction (5,6) for hybrid scanners, external tracking (7,8) or databased approaches (9–11) as well as combinations of different techniques. Methods that only make use of the data to identify and correct for motion are highly favorable because they are mostly independent of hybrid scanner set-ups or any additional equipment. Many data-based approaches focus on minimizing the difference between image frames, in terms of least squares of the residuals (12) and maximizing mutual information or voxelwise correlation, for example. Such data-based methods have been implemented in several preprocessing toolboxes, e.g.SPM12’s realignment (12).

As opposed to static or conventional dynamic PET, functional PET (fPET) uses a (bolus +) constant infusion protocol, providing enough activity to image different activation states or connectivity within only one measurement (13–16). Despite fPET being a rather novel technique, continuous technological advancement has made it a valuable tool to explore rapid metabolic dynamics within the brain. Specifically, dynamic and thus also fPET’s temporal resolution has gradually improved from 60s to 3s (13,16–18) per image frame and is now pushing the single and sub-second realm (19). Such high temporal resolution frames, however, generally exhibit a low signal-to-noise ratio (SNR). This is due to detector specifications as well as the nature of the decay that is used for signal generation in PET, as only a few counts are assigned to individual images. Furthermore, effects such as photon noise, detection of random events and scatter further diminish the data quality. Therefore, the performance of data-based motion correction of fPET data with short frame lengths is unclear. This is however relevant for the quantification of the brain’s physiological processes, such as energy metabolism. Thus, a re-evaluation of the adequacy of current techniques is necessary to ensure the integrity of research findings.

The inherently low-count frames present at the beginning of dynamic PET acquisitions have motivated the development of advanced post-processing methods to enhance image quality and facilitate data preparation and analysis, which can also be leveraged for high temporal fPET. These include methods based on the application of deep learning for image generation with simultaneous SNR enhancement (20,21). However, it remains unclear whether these methods provide substantial improvements in the further processing of the imaging data, specifically for motion correction.

In this work, we aim to evaluate the effect of two different SNR-enhancing algorithms on databased motion correction as compared to the default correction of the unmodified data. Individual fPET frames with a duration of 3s each were subject to a realignment algorithm for rigid motion following either i) no SNR enhancement, ii) SNR enhancement by application of a deep learning algorithm or iii) SNR enhancement by using a sliding window moving average.

## 2. Materials and Methods

### 2.1. Subjects and Study design

Nineteen healthy participants (age ∼ 22.9 ± 2.3 years, 7 male participants) underwent a single simultaneous PET/MR measurement. Each participant was examined prior to their study participation to ensure their eligibility. The examinations included routine blood parameters, neuropsychiatric assessments as well as electrocardiography. Exclusion criteria comprised pregnancy or breast-feeding, MRI contradictions, substance abuse or current treatment with medication, previous radiation exposure during a study and somatic/neurological/psychiatric pathologies. Participants were insured and received financial compensation for their time. The subjects provided written informed consent prior to their participation. All procedures were conducted in agreement with the Declaration of Helsinki and with approval of the Ethics Committee of the Medical University of Vienna (ID number 1479/2015).

Participants were subjected to a 20 min PET/MR measurement. The scan was separated in 8 min of baseline acquisition (fPET only acquisition) and 12 min of working memory task performance (simultaneous BOLD-fMRI and fPET acquisition). Cognitive task performance was arranged in a block design, including 10 x 36 s blocks and respective resting periods after each task block. The measurement was preceded by a T1-weighted structural MRI.

For cognitive stimulation, the n-back working memory task was selected. During task performance, a sequence of letters of the German alphabet was provided (b, c, d, g, p, t, w), with the letters being shown for half a second each. The task included two conditions: the 0-back control condition, which requires the identification of the letter “X”, and 2-back condition, during which the subjects needed to decide whether the current letter is the same as the second to last letter. These tasks had to be completed while disregarding upper/lower case. The two conditions were provided in a pseudo-randomized order and required a positive decision (“the letter fulfills the respective condition”) via a button press. Instructions for the current condition were shown at the beginning of each block. The baseline as well as the resting periods were accompanied by the display of a crosshair. Participants were instructed to fixate the crosshair with their gaze and let their thoughts wander freely. A test run for each condition was performed right before the measurement inside the scanner. Subjects had to use only their right index finger during the task.

### 2.2. Data Acquisition and Processing

fPET images were acquired using the Siemens Biograph mMR scanner. Structural T1-weighted images were attained using a MPRAGE sequence (TE/TR = 4.21/2200 ms, TI = 900 ms, flip angle = 9°, matrix size = 240 × 256, 160 slices, voxel size 1 × 1 × 1.1 mm, 7.72 min). Thereafter, fPET data were acquired, during which the radiotracer [^18^F]FDG was administered according to a bolus + constant infusion protocol (816ml/h for 1 min, 114.9ml/h for 19 min, 5.1 MBq/kg body weight, 20 min). The BOLD-fMRI was recorded as an EPI sequence (TE/TR = 30/2000 ms, flip angle = 90°, matrix size = 80 × 80, 34 slices, voxel size = 2.5 × 2.5 × 3.325 mm, 12 min).

Altogether, 15 arterial blood samples (every 20 s during the first 3 min, afterwards at 4, 5, 8, 12, 16, 20 min relative to the start) were drawn, partially using a VAMP Plus system (Edwards Lifesciences, applied after 5 min post infusion start) to minimize interference during the measurement. The blood samples were retrieved from the left radial artery. A gamma counter was used to determine whole-blood activity and plasma activity for the last six samples. Arterial input functions were retrieved by using whole-blood data and respective plasma-to-whole blood ratios. Prior to the measurements, blood glucose was determined as an average value of three individual, consecutive recordings to ensure fasting glucose levels of subjects.

List-mode data were reconstructed to 3s frames with each frame amounting to a matrix size of 344 x 344 and 127 slices (voxel size = 2.09 × 2.09 × 2.03 mm). Attenuation correction was performed using a pseudo-CT method based on the T1-weighted structural MRI (22). To ensure comparability between the different methods, the reconstructed images were later adapted to a matrix size of 128 x 128 and 128 slices. These are the dimensions previously proposed for the application of the selected deep learning algorithm (20,23) and to shorten the overall processing time.

Preprocessing was performed using SPM12 and included realignment with different strategies (see below), and spatial normalization to MNI-space, using the structural images. Subsequently, data were smoothed using an 8 mm Gaussian kernel and masked to include only gray matter voxels.

For quantification of the fPET data, a general linear model (GLM) was used, separating the baseline from the task activation (15,16,18). The model was defined by two task regressors, for the 0- and 2-back condition, respectively, one regressor accounting for motion and one baseline regressor. The task regressors were modelled by ramp functions with a slope of 1 kBq/frame during task performance and 0 kBq/frame otherwise. The baseline regressor was defined as the average TAC across the gray matter, excluding those voxels that were identified as active during the analysis of the BOLD fMRI data. A principal component analysis of realignment parameters was used to reduce dimensionality, where the number of components were determined using the elbow method. Finally, the respective predictors for task activation as obtained from the GLM and the arterial input function were used as input for Patlak plots (24). The net influx constant K_i_ was determined as the slope of the plot after linearity had been established (linear fit from approximately 11 min after infusion start). The cerebral metabolic rate of glucose was not determined due to missing blood glucose levels of a single participant.

Extraction of the BOLD signal and processing of fMRI data were performed in SPM12 as in our previous work (18,25). Preprocessing included slice time correction and realignment (quality = 1, registration to the mean frame). Further processing was not required, in this case, as only the realignment parameters were of interest for the further analyses.

### 2.3. fPET Motion Correction Strategies

All motion correction procedures were carried out with SPM12’s realignment (quality = 1, registered to overall mean after prior registration to mean of motion-free sequence). In addition to the unprocessed data (i.e., standard SPM-based motion correction), we tested two additional approaches for the improvement of the SNR prior to realignment. Such an increase in SNR is hypothesized to lead to a better realignment. Importantly, the modified fPET data with increased SNR were only used for the estimation of motion parameters, but not for the subsequent quantification. Thus, individual sets of motion parameters were retrieved which were then used for the correction of the original, unmodified data (see fig. 1). In the following, “standard approach” or similar expressions generally refer to the realignment without application of any prior SNR enhancement techniques, although all of the mentioned techniques are subjected to the same realignment procedure after possible pre-treatment of the data.

**Figure 1:**
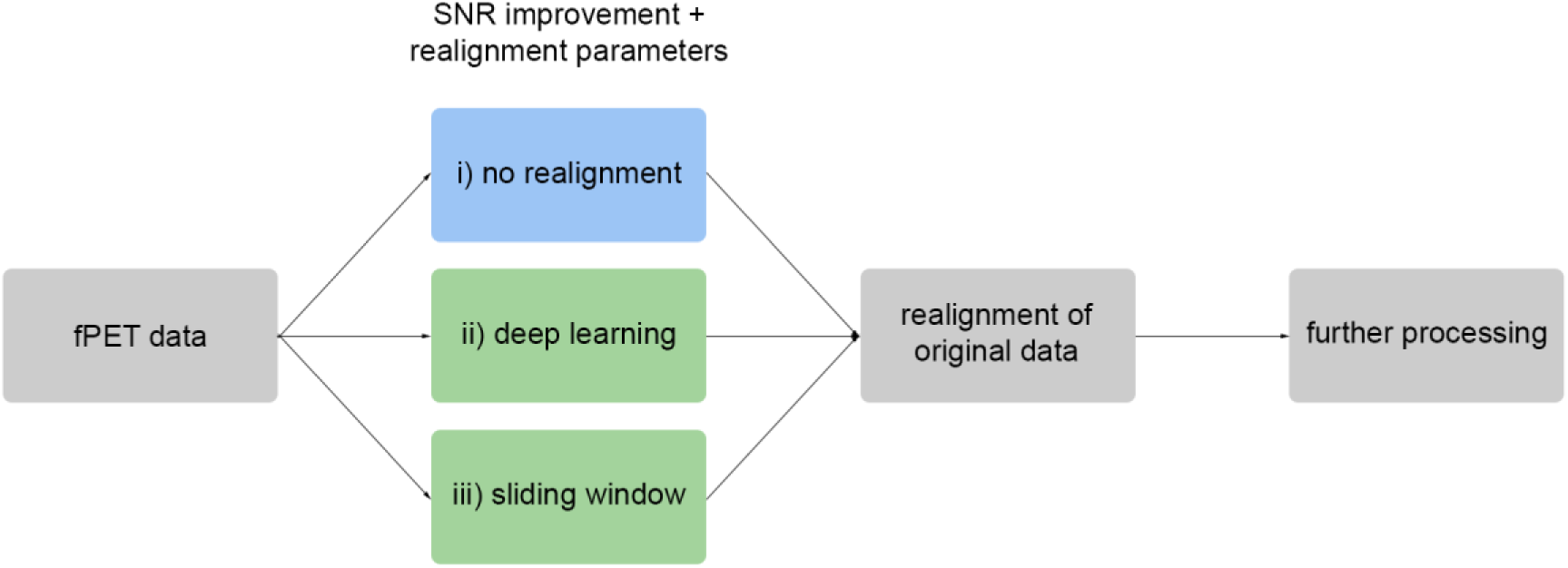
Visual representation of the processing strategy. The three separate approaches are employed for motion correction: i) without prior processing of the images, ii) using a deep learning algorithm or iii) application of a sliding window moving average. Approaches ii) and iii) are applied only to increase the SNR of fPET images and retrieve motion parameters that are subsequently used for the realignment of the original data. Further processing includes spatial normalization as well as quantification and statistical comparisons.

#### Deep-learning-based SNR Enhancement

The deep-learning-based SNR improvement was implemented by using a previously published software, PET nAvigators usiNg Deep leArning (PANDA) (20,23). The software uses a conditional generative adversarial network for generation of artificial activity patterns that can thereinafter be used for motion parameter retrieval. The algorithm is based on a mapping strategy that uses the pairing of early image frames, which generally display a low SNR and thus complicate motion correction, and high-SNR late frames. For the data at hand, a volume-to-volume mapping was created by the algorithm using a training set (13 subjects), and then subsequently applied to a test set (remaining 6 subjects) to improve the SNR of these image frames, specifically. As the sample size is quite limited with 19 subjects in total, it was crucial to be able to use all of the subjects for further analyses. Thus, training and test sets were alternated in four sequential iterations, to achieve the generation of one artificial series of volumes per subject included in the dataset.

To enable the mapping procedure, a reference for a high-SNR frame had to be selected. This was determined to be the mean of the last full minute of acquisition (20 frames). Theoretically, a mapping (and thus a corresponding training) would need to be conducted for each frame of the time series. However, since the time series contains more than 380 3s-frames for each of the subjects, this was not feasible because mapping of 380 frames would take approximately 380 times 7 hours per subject. Therefore, a corresponding “mapping frame” was determined as follows: The summed voxel intensities were computed for each frame of a sequence. The first frame whose summed voxel intensities exceeded 10 MBq (around 3-4 minutes after infusion start) was selected. To this time point three minutes were added and the resulting frame was then chosen as the respective “mapping frame”. This was done to ensure that the mapping frame would contain sufficient signal strength to be able to provide a successful mapping. For each subject in the dataset, the matched pair of these two frames was used in the training process. To ensure that the selected mapping frame is representative for the individual time courses, mutual information content was analyzed to identify potential singularities. Once the mapping was completed, the SNR of each frame was increased by applying it to the dynamic data, respectively.

#### Sliding-window-based SNR Enhancement

For the sliding window SNR enhancement, a moving weighted average of 5 frames at a time (15s of acquisition) was used to replace the original frames. To retain the general activity patterns observed, the weighting function that was selected for calculating the average was an exponential function (26) to the power of 0.5, which assigns the present frame in the sequence a contribution of 1, and the four previous frames correspondingly smaller weights. The following formula (1) describes the time-dependent weights, with n assuming values from 1 to N = 5, indicating the respective frames that are being averaged:

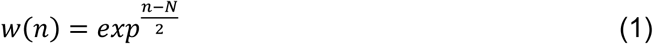

### 2.4. Performance Evaluation and Statistical Analysis

#### Comparison of SNR

A general assessment of the SNR improvement for each approach was carried out. Furthermore, the mean normalized mutual information per frame for each of the three separate techniques was analyzed (20), calculated with regard to the reference as used for the deep learning algorithm, i.e., the mean of the last full minute of acquisition. We also analyzed voxelwise Pearson’s correlation between each frame and said reference, as well as integrated voxel intensities, to assess differences in time-dependent signal strength, and the SNR in terms of activity distribution mean in relation to standard deviation. A custom mask was used, which was calculated by thresholding the reference volume to only include values > 1 kBq.

#### Comparison of motion parameters

Realignment parameters obtained from the fPET methods were compared among each other and with the BOLD fMRI reference, using correlations between parameters over time, z-transformed and averaged across six degrees of freedom. BOLD fMRI data were used as a reference standard for comparison as they were acquired simultaneously during task performance (6,27). The fPET sequences were cropped to coincide with the interpolated fMRI sequences.

Finally, to quantify the divergence between the methods, the sum of the absolute differences between the respective fPET and fMRI realignment parameters across time was compared. To retrieve a collective measure including different degrees of freedom, summed absolute differences of framewise displacements between fPET and fMRI were also calculated.

#### Comparison of activation

As the quality of motion correction can substantially affect the outcome, we also compared the task-specific changes in glucose metabolism for both, control and task conditions, retrieved by application of the GLM and subsequent quantification. We did so on a group level, performing one-sample (task > baseline) and, for inter-method comparison, paired t-tests, each subsequently thresholded with p < 0.05 at the FWE-cluster-corrected level, after p < 0.001 uncorrected voxel-wise threshold.

## 3. Results

### Comparison of signal-to-noise ratios

When comparing the signal characteristics of the individual techniques to the unmodified data, we observed substantial improvements in image quality, as quantified by signal strength and/or noisiness, for both SNR enhancement methods (see fig. 2). These improvements were especially pronounced for the application of the deep learning algorithm (fig. 2K-O). However, it is also apparent that the structural integrity of (sub-)cortical areas might not be preserved by the algorithm, as visible in the thalamus for example, although anatomic distributions are difficult to evaluate using only PET images. Another observation with regard to this algorithm specifically, is the constant and inflated signal strength, independent of time passed since radiotracer injection start (suppl. fig. 1C). For the sliding window technique, we observed slightly lower signals as compared to the standard approach, as induced by the weighting function, however, the noise content was apparently reduced as well (fig. 2P-T, suppl. fig. 1C-D). To better illustrate the practical relevance of these results, another time series was assessed: the original unmodified fPET frames, which were subjected to smoothing using a 5 mm Gaussian kernel. This internal smoothing procedure of SPM also produces a slight reduction of the noise content (fig. 2F-J).

**Figure 2:**
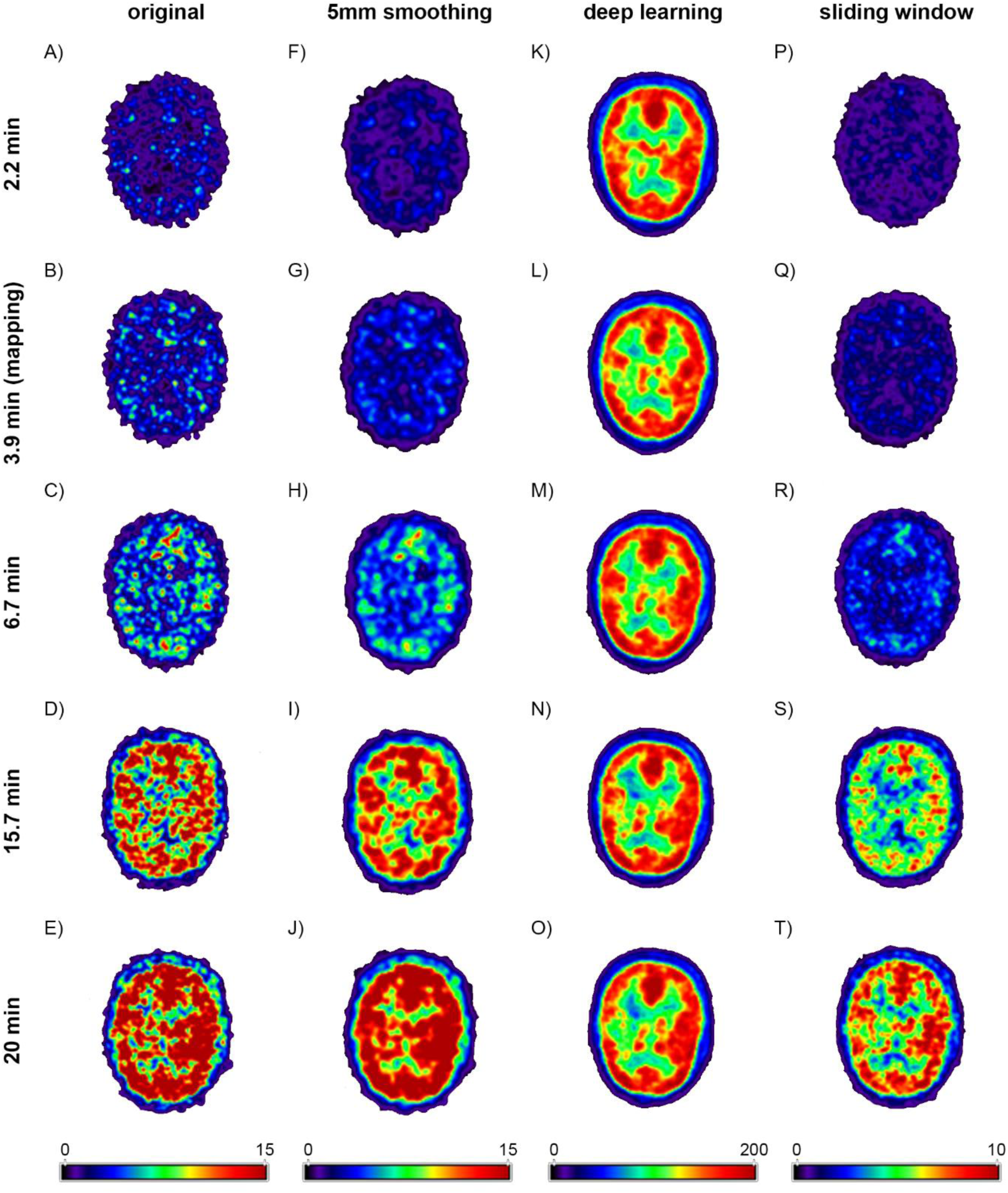
Overview of the image quality at different time points for the separate SNR-modifications performed. The different procedures include the deep-learning-based method and sliding window technique, as used for extraction of the motion parameters hereinafter. Furthermore, besides the original data, smoothed images (5 mm Gaussian kernel, i.e., SPM12’s internal smoothing) are displayed as well. The time stamps refer to the time passed relative to the start of administration of the radiotracer. The slices used for comparison were extracted at z = 18 mm MNI-space. The colorbars illustrate the different signal strengths in kBq/cm³, respectively. The absolute values corresponding to the sliding window approach are reduced due to the structure of the weighting function, which is normalized to assign a value of one to the frame to be corrected and respectively smaller values to the previous four frames.

Analysis of mean mutual information and voxelwise correlation between individual frames and reference frames showed distinct increases of both quantities for both SNR-enhancing methods (suppl. fig. 1A-B). Furthermore, the courses of mutual information for the individual subjects were used to ensure that the selected mapping frames did not suffer from any anomalous signal characteristics that might deteriorate the overall mapping quality. Regarding the integrated voxel intensities over time, the deep learning and sliding window techniques showed a distinct reduction of noisiness. Finally, for the direct comparison of the average SNR we found that the deep-learning-based technique dominates for the early frames of the sequence, but declines over time (suppl. fig. 1D). Eventually, the sliding window approach yields the best results for later frames, however, all three methods perform more similarly as the measurement advances in time. A single dataset was excluded from these calculations due to a significantly delayed start of the tracer application.

### Comparison of motion parameters

Across the three methods, the sliding window technique shows the overall highest correlation with the fMRI motion parameters (r = 0.808 ± 0.381, see fig. 3A). The standard realignment achieved slightly though significantly higher correlations (r = 0.652 ± 0.304) compared to the deep learning implementation (r = 0.547 ± 0.239). These observations generally hold also for individual subjects (see suppl. fig. 2). The three realignment techniques revealed the highest correlations for the comparison of the standard realignment and the sliding-window-based approach (fig. 3B), indicating a high similarity of fPET motion parameters (r = 0.904 ± 0.186).

**Figure 3:**
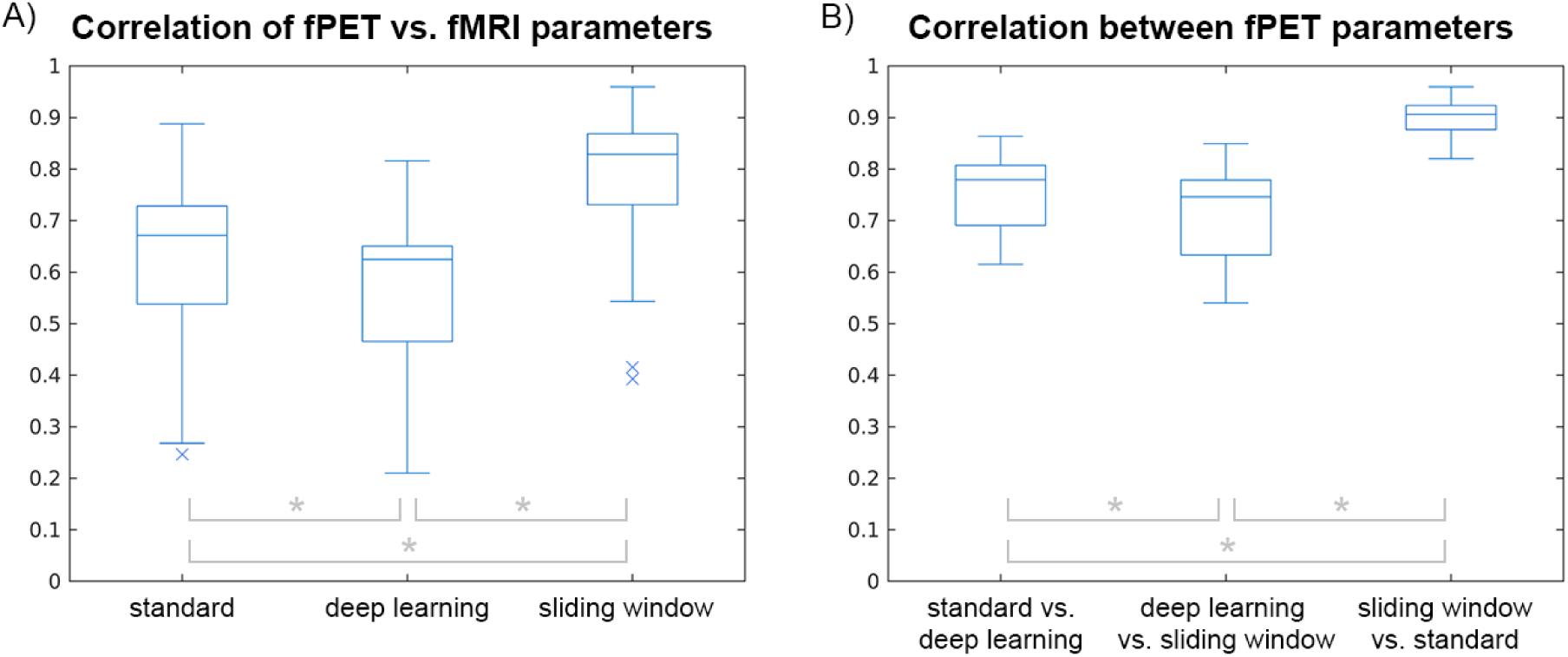
Average correlations of motion parameters: (A) Correlations between motion parameters as obtained from fPET data vs. those obtained with fMRI. (B) Correlations of the three motion correction methods with each other for fPET data. The mean sequences were obtained by combining the correlations of each of the six individual parameter courses after a z-transformation to account for divergence from normal distributions. Box plots show data across all subjects, * p < 0.001 with paired t-tests.

Finally, time courses of motion parameters show that the motion correction techniques are prone to larger shifts that potentially overcorrect as compared to fMRI parameter sequences (see suppl. fig. 3, table 1). However, of the three techniques, the sliding window appears to smooth out such instantaneous and high changes to a higher extent (suppl. fig. 3, yellow time course). To assess the divergence of parameters further, we calculated the differences in framewise displacement as a consolidated measure for all individual motion parameters. Here, we observe a distinct similarity of the sliding window method, with respect to minimizing the absolute difference compared to the fMRI reference (see table 1, suppl. table 1), which is also apparent from the exemplary time courses (suppl. fig. 3). On the level of individual motion parameters, we compared the sum of the absolute difference between the parameters’ time courses (fPET and fMRI). Differences in translations are of a similar extent for the standard realignment and sliding window approach. Considering rotational degrees of freedom, values were in a similar range on the individual-parameter level (table 1).

**Table 1:** Absolute summed differences in framewise displacement (FD), translations and rotations, compared between fPET and fMRI data (average across subjects, standard deviations displayed in brackets) and calculated for each realignment method, separately. The sum encompasses the absolute differences at each time point of the simultaneous measurement, with respective offset-correction for the fPET data.

| Outcome param. | Standard realignment |  |  | Deep learning |  |  | Sliding window |  |  |
| --- | --- | --- | --- | --- | --- | --- | --- | --- | --- |
|  | x | y | z | x | y | z | x | y | z |
| $\Delta FD$ [mm] | | 73.92<br>(16.45) | | | 112.90<br>(15.83) | | | 18.72<br>(4.97) | |
| $\Delta Transl.$ [mm] | 38.05<br>(16.54) | 56.34<br>(23.77) | 41.32<br>(23.10) | 45.38<br>(17.17) | 67.48<br>(36.74) | 67.38<br>(33.25) | 24.75<br>(12.85) | 50.98<br>(24.78) | 32.64<br>(22.89) |
| $\Delta Rot.$ [°] | 1.15<br>(0.49) | 1.01<br>(0.57) | 1.17<br>(0.56) | 1.22<br>(0.59) | 1.14<br>(0.48) | 1.22<br>(0.39) | 1.13<br>(0.57) | 0.72<br>(0.37) | 0.59<br>(0.32) |

### Comparison of activation

Similar to our previous work (18), we identified significant activation in regions typically addressed during working memory challenges such as the n-back task (28,29). Qualitatively, the comparison of the different activation patterns revealed high similarity between the three methodologies (see fig. 4).

**Figure 4:**
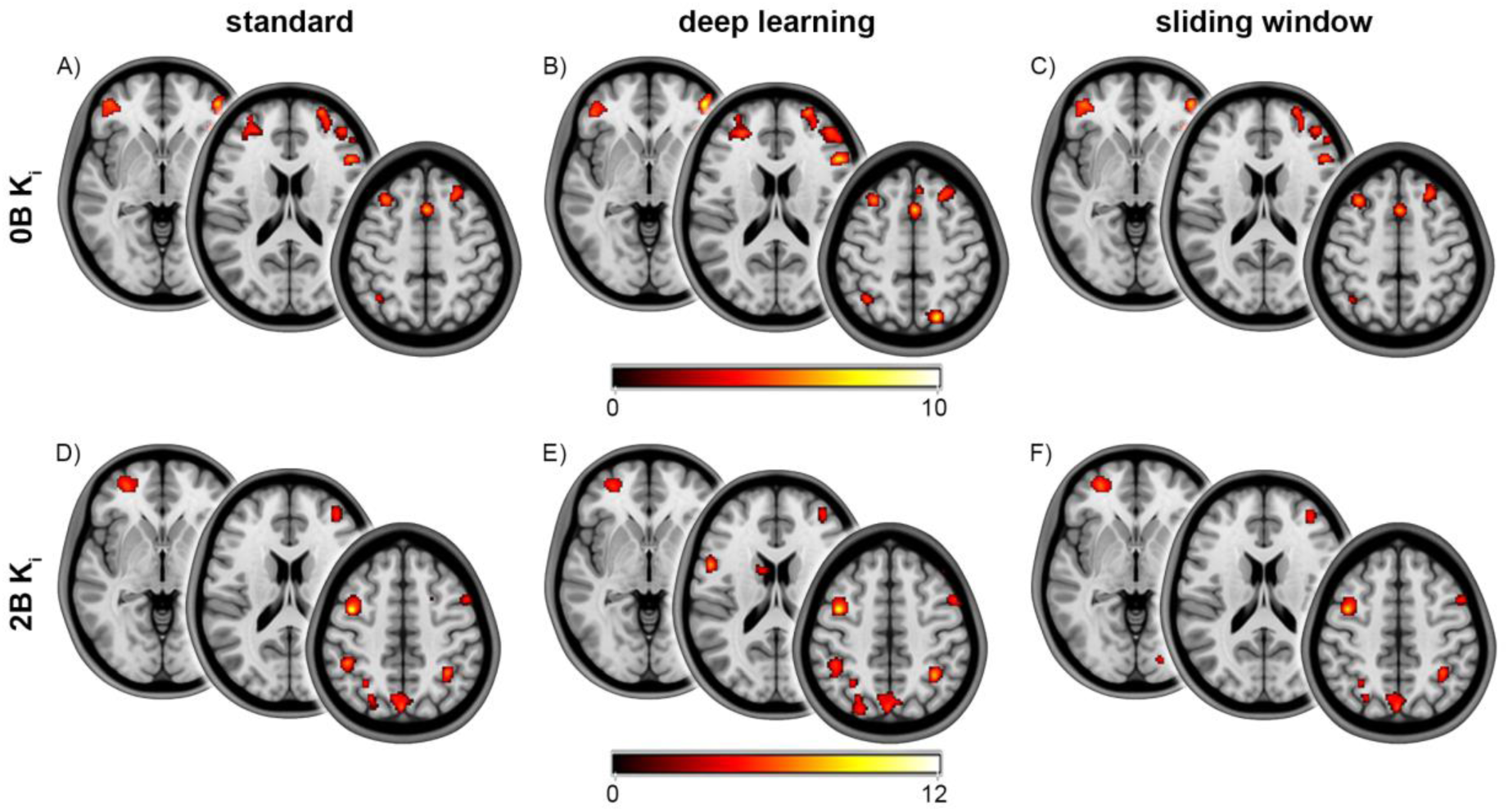
One-sample t-maps displaying the significant activation for both task conditions, 0-back (A-C) and 2-back (D-F), and the three different motion correction techniques with and without SNR enhancement. The maps reveal the significant clusters of activation for the standard realignment (A,D) as well as the deep-learning (B,E) and sliding-window-based SNR enhancement (C,F) prior to the realignment of the dynamic frames. Slices were extracted at z = -3 / 18 / 52 mm (A-C) or z = -3 / 18 / 48 mm (D-F) MNI space (left to right). These slices were selected as they exhibit the greatest overall changes in cluster differences between approaches. The colorbars indicate the range of t-scores (p<0.05 FWE cluster corrected, following p<0.001 voxel uncorrected).

To further assess whether there are quantitative differences in the identification of task effects between the different approaches, paired t-tests were used. The analyses revealed no significant differences between the sliding window and standard method, for neither of the task conditions. However, for the comparison of the deep-learning-based technique and the standard approach, significant differences were identified for both task conditions (see fig. 5). These significant clusters (deep learning < standard) were mainly located in the occipital lobe or around parts of the transversal/sagittal sinus but did not concern any of the regions with task-specific changes in metabolism (28,29) (fig. 5).

**Figure 5:**
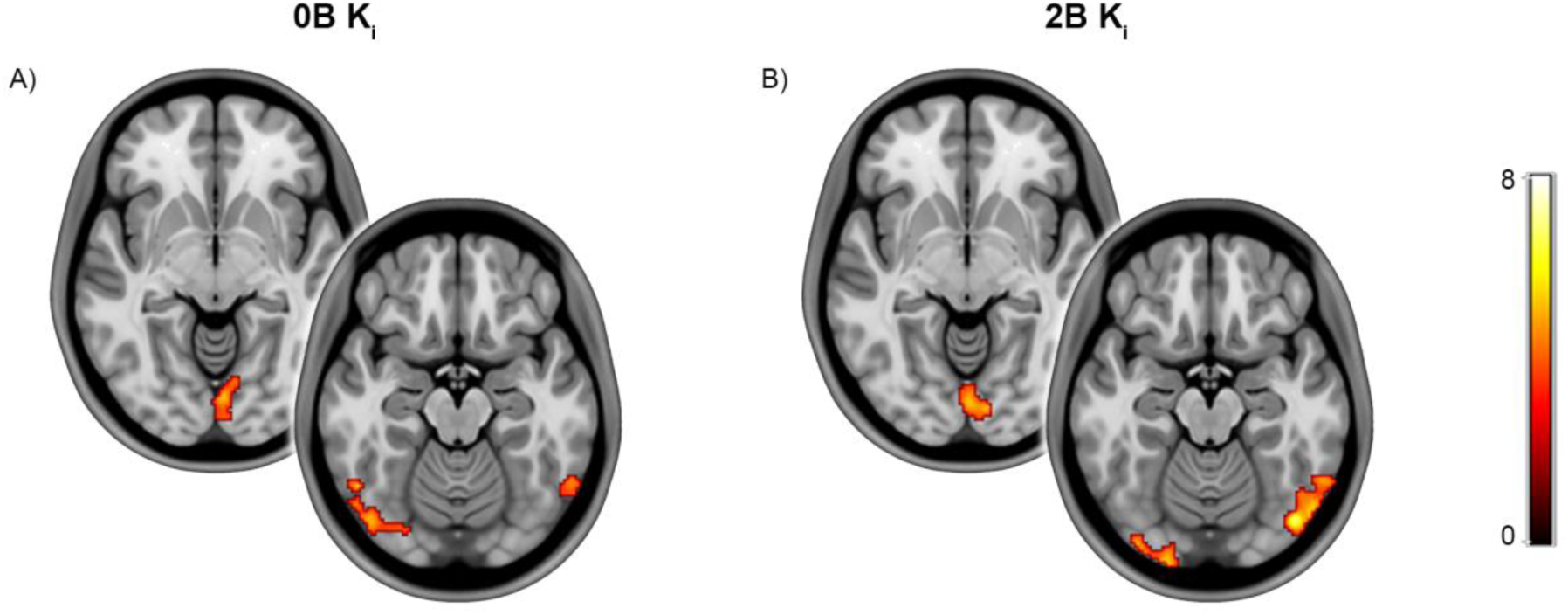
Results of the paired t-tests comparing the deep-learning approach as compared to standard realignment, for both 0-back (A) and 2-back (B) conditions, deep learning < standard realignment. The t-maps showed significant differences only in regions that did not exhibit activation in the previous one-sample t-tests (fig. 4). The slices were extracted at z = -7 mm (left) and z = -17 mm (right) MNI space and selected as they included the majority of significant clusters. The colorbar indicates the range of t-scores (p<0.05 FWE cluster corrected, following p<0.001 voxel uncorrected).

## 4. Discussion

We assessed whether motion correction of high temporal resolution fPET data benefits from prior SNR enhancement as compared to standard realignment. Although the deep learning and sliding window algorithms artificially increased the SNR of fPET data, improvements in motion correction were limited. The sliding window approach showed highest similarity with motion parameters obtained from simultaneously acquired fMRI data, followed by standard realignment and the deep learning approach. Comparison of neuronal activation patterns revealed that there was high similarity in task-specific clusters between the methods, whereas quantitative differences only occurred for the deep-learning algorithm in unspecific brain regions. Altogether, these results do not support the hypothesis that such approaches providing an enhancement of SNR inflict an improvement compared to established motion correction.

The deep-learning-based technique led to distinct improvements regarding signal strength and noise content of the individual volumes. With respect to the comparison of motion parameters, however, we found that this method performed worse or at most comparably to the standard approach. Notably, the modification of the original data introduced extensive shifts as compared to the other techniques, potentially exceeding reasonable bounds. This overmodulation could also be one of the reasons for the apparent lack of improvement, especially with regard to task activation patterns. The comparison of glucose metabolism did not show any substantial differences between the methods in the regions of interest for the respective task. Significant decreases compared to the standard approach were only observed in areas of the occipital cortex, which were however below the detection threshold of task effects. Furthermore, these differences were located around large blood vessels in the brain, i.e., the sagittal and transversal sinus. We speculate that the deep-learning-based SNR improvement might be more sensitive to motion artefacts introduced in these areas and especially emerging from blood vessels. The decreased sensitivity to motion-related artifacts may parallel previous work showing improved definition of image-derived input functions when using prior signal enhancement with deep learning (20). Although unexpected, the relevance for task analyses remains limited. Here, the lack of improvement might be indicative of more general shortcomings of the training and mapping routine. One of these shortcomings is its apparent failure to reproduce some of the anatomical structures and tendency to enhance potential artefacts, thus potentially impacting image registration.

The inter-individual variation of the output’s quality further highlights one of the more general struggles associated with the application of deep learning, namely the “black box” problem, i.e., the user not being privy to the intrinsic criteria and features used for training as well as issues regarding the output’s interpretability (30). Furthermore, the applicability and generalizability of the resulting mappings is also highly dependent on the quality and amount of input data (31,32), the latter being a limitation of this and most other neuroimaging studies. Nevertheless, deep learning could still provide a possible solution for the appropriate mitigation of movement artefacts (21,33) in future applications, especially when refining the model inputs and applying respective methods to overcome limited data availability and challenges of generalizability, explainability and interpretability (30,32,34).

A more distinct difference in performance with regard to the motion parameters was found for the sliding window SNR enhancement, which showed highest similarity with the fMRI-based reference. However, for neuronal activation patterns, the approach did not show any significant differences compared to the standard approach. This lack of change and the improvement in terms of motion parameters appear to be conflicting. However, the match of motion parameters can be largely attributed to the increased temporal smoothing that occurs when applying a sliding window averaging to the image sequence, which reduces overall divergences. This smoothing is also apparent, when considering the integrated voxel intensities, where the average noise content across time is reduced drastically.

Other important considerations which limit the usage of deep learning for processing high temporal resolution data specifically, are computational power and overall efficacy. One of the four individually employed mappings took approximately 7.5 hours using a dedicated graphical pro-cessing unit, also depending on the frame dimensions (20). By reducing the number of mappings that has to be performed, we managed to decrease the overall processing duration. However, it is possible that the results we obtained might be affected by this simplified implementation, in the course of which we created only one representative mapping instead of roughly 380 for the entire dynamic sequence. The computation for the sliding window was naturally less complex, but still presents an additional effort of about 2.5 hours for the whole dataset as compared to the use of original data.

In summary, based on the comparison of motion parameters and neuronal activation patterns, along with considerations regarding implementation and processing time, it appears that realignment without prior SNR modification is sufficient. It should be emphasized, nonetheless, that this is true for the experimental conditions employed in this case specifically, i.e., for data with high temporal resolutions and corresponding low SNR values.

Our analyses are subject to several methodological limitations. For the deep-learning-based approach the sample size of the study population was small, which might introduce overfitting (32). This could result in the creation of suboptimal mappings and thus artificial volumes. Data augmentation, for example by using geometric transformations such as translations or rotations, can only resolve this partially (32). Still, the current sample size is similar to previous work (20) and the applied mappings were obtained by training with different subjects and iterating though the dataset. Another aspect to consider is that the deep learning application requires the pre-definition of several parameters to specify the network operation and performance. The precise alignment of these parameters could significantly affect the outcome with regard to the mapping quality, which however is again a time-consuming endeavor.

The effect of the sliding window technique is also defined by its parameters, such as window size and weighting functions. Modification of these might change these results, but need to be tested in future work. For instance, application of different weighting functions could potentially result in further performance improvements.

We chose the BOLD-fMRI parameters as a reference standard for the motion parameter comparison as their eligibility for PET motion correction has previously been established (6,27). Still, movement information retained from fMRI sequences might not capture the real extent of the motion as compared to external tracking devices (35). Finally, both methods might also perform better when applied in concordance with a different preprocessing routine. However, the SPM pipeline used in this work includes standardized parameters performed with a widely-used toolbox.

Despite the standard approach performing sufficiently for the data at hand, new methods might be required in future analyses as the temporal resolution is improved even further. The possibility of high temporal resolution imaging has created a manifold of new and re-introduced wellknown challenges. However, it has also opened up a realm of quickly growing research fields, such as rapid dynamics of cerebral metabolism (36) or comprehensive inter-modality functional imaging studies (18). In this context, the appropriate correction of motion artefacts could prove especially relevant. Among others, this includes the computation of metabolic and molecular connectivity (37,38). Analogous to the influence of motion on functional connectivity in fMRI (39,40), movement can distort the estimated interaction between brain regions, which might complicate the interpretation of coordinated inter-regional metabolic activity and the respective network-based organization. Thus, the question of suitable motion correction remains crucial, for applications in emerging fields of fPET imaging specifically.

## Conclusions

This study evaluated the usage of two separate methodologies for SNR enhancement prior to motion correction, specifically for high temporal resolution fPET data. Weighting the modest differences for the identification of neuronal activation against the computational cost, we demonstrate that the default realignment procedure without any prior SNR enhancement provides robust results. As current research efforts point towards the acquisition of even shorter dynamic image frames, the implementation of alternative methods based on different deep learning strategies and network architectures can still offer promising solutions for future advancements in the field of motion correction for neuroimaging data.

## Further statements

### Author Contributions

Conceptualization: A.H., M.R., P.F., Methodology: P.F., A.H., Writing – original draft: P.F., Writing – review and editing: all authors, Funding acquisition: A.H., R.L. All authors discussed the implications, revised the manuscript and approved the final version.

## Acknowledgements

We thank the graduated team members and the diploma students of the Neuroimaging Lab (NIL, head: R. Lanzenberger) as well as the clinical colleagues from the Department of Psychiatry and Psychotherapy for clinical and/or administrative support. In detail, we would like to thank S. Kasper, K. Papageorgiou, L. Silberbauer, C. Schmidt, B. Eggersdorfer, J. Unterholzner and V. Popper for medical support, L. Rischka for acquisition and analysis support, V. Ritter and C. Wotawa for subject recruitment. We are further grateful to W. Wadsak, V. Pichler, G. Karanikas, W. Langsteger and the radioligand synthesis team from the Department of Biomedical Imaging and Imageguided Therapy, Division of Nuclear Medicine for acquisition support and supervision. The scientific project was performed with the support of the Research Platform for Medical Imaging (RPMI) of the Medical University of Vienna.

## Data and Code availability

Raw data will not be publicly available due to data protection reasons. Processed data and custom code can be obtained from the corresponding author with a data-sharing agreement, approved by the departments of legal affairs and data clearing of the Medical University of Vienna.

