## Supplementary Material for "Evaluation of Data-based Motion Correction Techniques for High Temporal Resolution Functional PET"

For submission to

**bioRxiv**

#### Supplementary figures

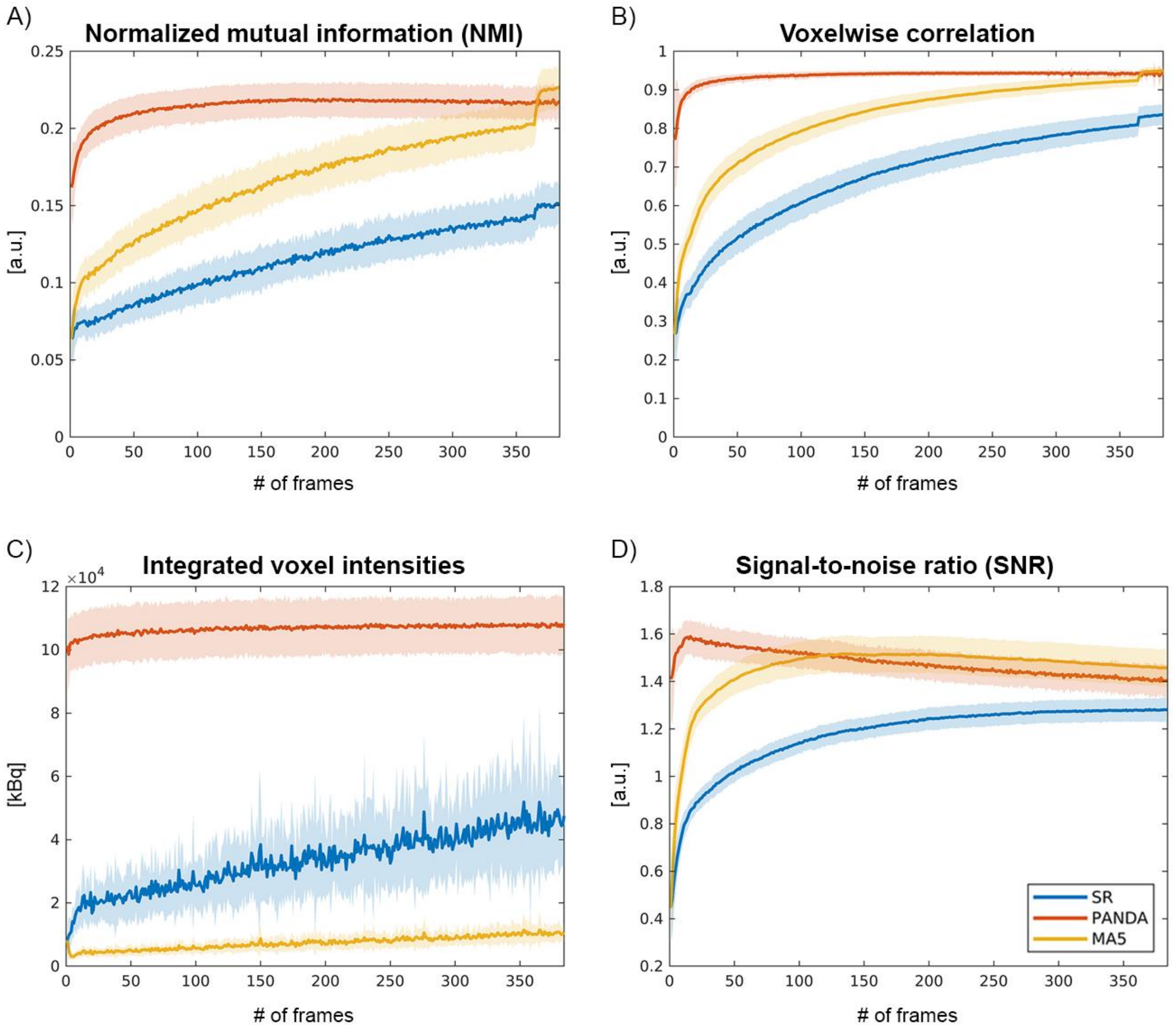


*Supplementary figure 1: Comparison of the different SNR-enhancement methods with respect to image quality and noise. Mean normalized mutual information (NMI, A) and voxelwise correlation of time frames (B) were calculated with respect to the reference frame that encompasses the last full minute of acquisition (i.e., mapping frame of the deep-learning algorithm). Integrated voxel intensities (C) and simplified SNR (D) was compared between the modalities across time. The shaded areas present the respective standard deviations calculated across subjects. The abbreviations in the legend refer to standard realignment (SR), deep learning based realignment (PANDA) and the weighted sliding window average of five frames (MA5).*

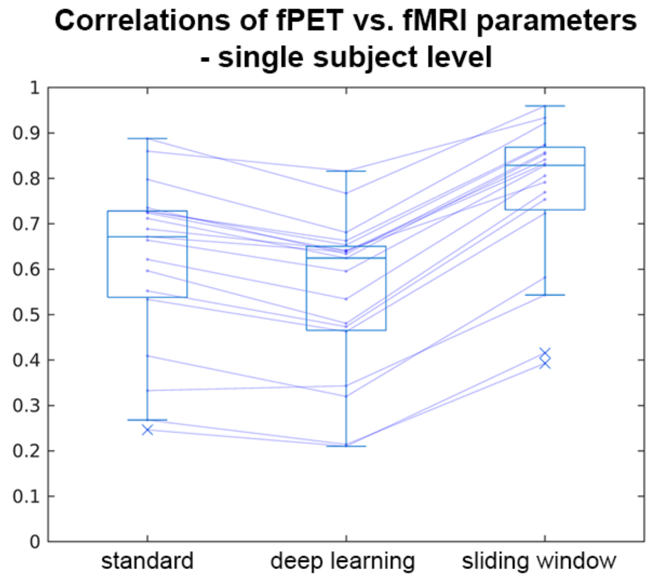


*Supplementary figure 2: Correlations between PET-based and fMRI-based motion parameters on a single subject level (each line is one subject).*

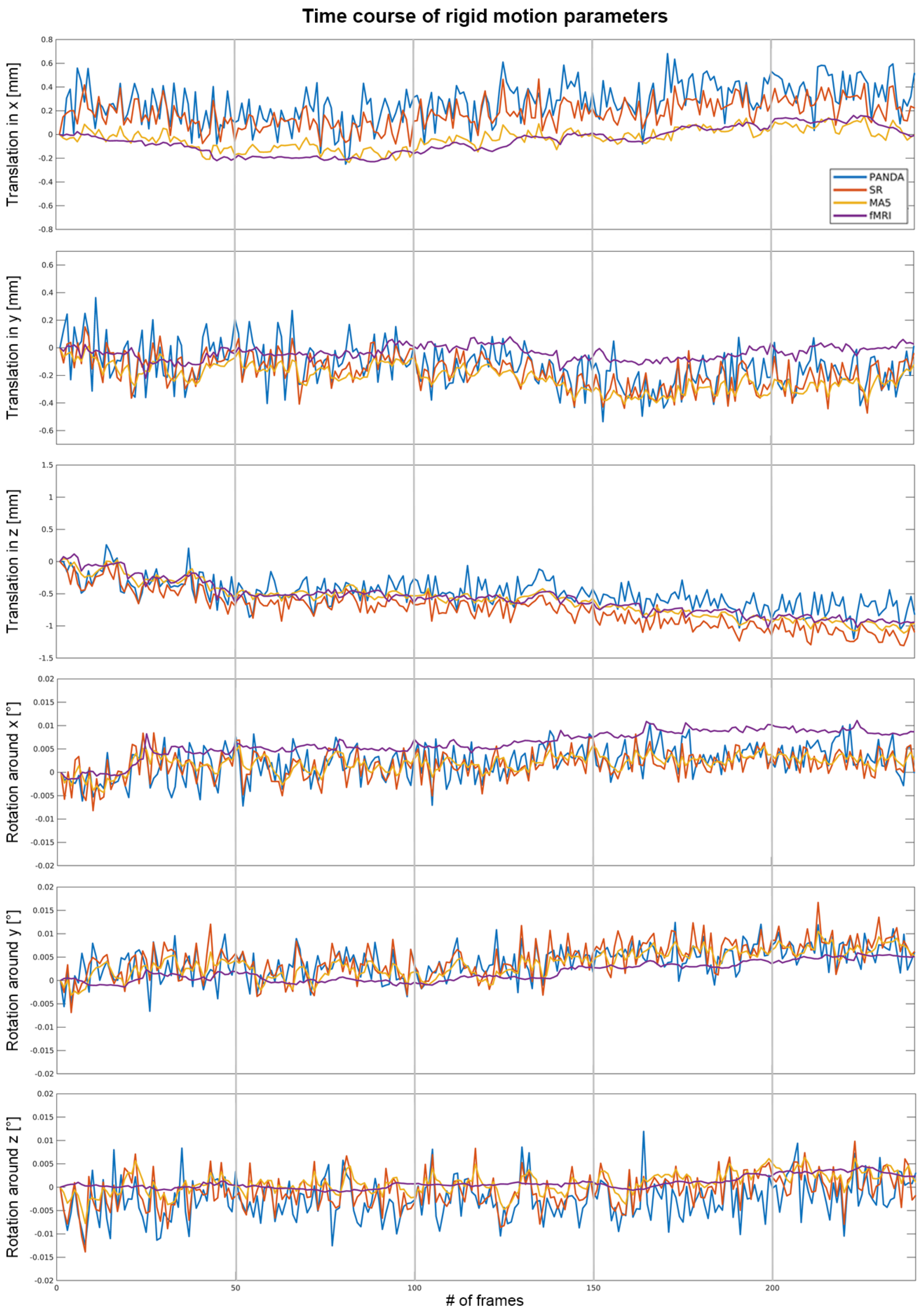

Time course of rigid motion parameters
Translation in x [mm]
Translation in y [mm]
Translation in z [mm]
Rotation around x [°]
Rotation around y [°]
Rotation around z [°]
PANDA
SR
MAS
fMRI
### of frames

*Supplementary figure 3: Exemplary time courses of the motion parameters for the different realignment approaches of a representative subject. They illustrate the behavior of the translational and rotational degrees of freedom, separately, and span the period of simultaneous PET/MR acquisition (~240 frames). The abbreviations in the legend refer to standard realignment (SR), deep learning based realignment (PANDA) and the weighted sliding window average of five frames (MA5) as well as the fMRI realignment parameters.*

#### Supplementary tables

| Outcome param. | Standard realignment | | | Deep learning | | | Sliding window | | | BOLD fMRI | | |
|---|---|---|---|---|---|---|---|---|---|---|---|---|
| | x | y | z | x | y | z | x | y | z | x | y | z |
| FD [mm] | | 93.54 (14.56) | | | 133.05 (14.38) | | | 32.43 (4.75) | | | 20.73 (4.72) | |
| Transl. [mm] | 62.48 (23.00) | 63.19 (30.77) | 80.47 (57.93) | 79.23 (30.31) | 63.86 (30.59) | 111.31 (67.37) | 51.78 (25.54) | 61.88 (37.38) | 74.98 (52.71) | 61.04 (35.10) | 55.34 (48.82) | 88.03 (68.34) |
| Rot. [°] | 1.77 (1.01) | 1.58 (0.70) | 1.53 (0.80) | 1.42 (0.96) | 1.64 (0.76) | 1.21 (0.49) | 1.76 (1.05) | 1.21 (0.59) | 0.99 (0.54) | 1.58 (1.24) | 1.15 (0.77) | 1.03 (0.62) |

*Supplementary table 1: Absolute summed values of framewise displacement (FD), translations and rotations (average across subjects, standard deviations displayed in brackets) as calculated for each realignment method for fPET and fMRI data, separately. The sum encompasses the absolute values at each time point of the simultaneous measurement, with respective offset-correction for the fPET data.*
